# Mitochondrial phylogeography of grassland caterpillars (Lepidoptera: Lymantriinae: *Gynaephora*) endemic to the Qinghai-Tibetan Plateau

**DOI:** 10.1101/2023.03.08.531811

**Authors:** Ming-Long Yuan, Ming-Hui Bao, Qi-Lin Zhang, Zhong-Long Guo, Min Li, Juan Wang

**Author notes:** Correspondence and requests for materials should be addressed to M.L.Y. These authors contributed equally to this work (Ming-Long Yuan, Ming-Hui Bao, Qi-Lin Zhang).

## Abstract

Grassland caterpillars (Lepidoptera: Lymantriinae: *Gynaephora*) are the most damaging pests to alpine meadows in the Qinghai-Tibetan Plateau (QTP). Here, we conducted extensive sampling from 39 geographic populations covering almost the entire distribution of the eight *Gynaephora* species in the QTP to investigate phylogeographic patterns and speciation based on two mitochondrial genes (*cox1* and *nad5*). A total of 40 haplotypes were detected in the 39 populations, with >70% of haplotypes specific to single population. The monophyletic QTP *Gynaephora* migrated from non-QTP regions during the Pliocene, corresponding to the uplift of the QTP, suggesting a mode of transport into the QTP. Among the eight QTP *Gynaephora* species described by morphological characteristics, two species (*G. alpherakii* and *G. menyuanensis*) were recovered as monophyletic groups (Clades B and C), while the remaining six species formed two monophyletic clades: Clade A (*G. qinghaiensis*, *G. jiuzhiensis*, and *G. qumalaiensis*) and Clade D (*G. aureate*, *G. rouergensis*, and *G. minora*). These results suggested that the number of the QTP *Gynaephora* species may be overestimated and further studies based on both morphological and nuclear gene data are needed. Genetic differentiation and speciation were driven by intensive uplifts of the QTP and associated climate fluctuations during the Pleistocene, suggesting that isolation and subsequent divergence was the dominant mode of speciation. The Sanjiangyuan region (i.e., Clade A, characterized by high genetic diversity) may have been a glacial refugium of the QTP *Gynaephora*, as supported by analyses of gene flow and biogeography. High levels of genetic diversity were found in QTP *Gynaephora,* without population expansion, which may explain the high-altitude adaptation and outbreaks of grassland caterpillars in alpine meadows of the QTP. This study provides the largest phylogeographic analysis of QTP *Gynaephora* and improves our understanding of the diversity and speciation of QTP insects.

## Introduction

The Qinghai-Tibetan Plateau (QTP) has the highest average altitude in the world and is characterized by extreme environmental features (e.g., hypoxia, cold climate, and high levels of ultraviolet radiation). Extensive uplifts of the QTP since the Miocene period (∼23 million years ago, Ma) have caused dramatic climatic and environmental shifts (Harrison et al., 1992;Wu et al., 2001;Jia et al., 2003). Both geological and climatic changes are well-established drivers of species diversification and speciation patterns in various QTP plants and animals (Lei et al., 2014;Favre et al., 2015;Xing and Ree, 2017). Thus, the QTP and adjacent regions have numerous endemic species and have been recognized as biodiversity hotspots (Myers et al., 2000).

During the past few decades, the QTP has been considered a “natural laboratory” for exploring adaptation and evolution (Jiang et al., 2018). Phylogeographic analyses of many QTP taxa, including mammals (Zhang et al., 2005;Liu et al., 2012), birds (Zhang and Fritsch, 2010), amphibians (Che et al., 2010;Zhou et al., 2012;Yan et al., 2013), and fishes (Chen et al., 2004; Peng et al., 2006; Qi et al., 2006), have been reported. An increasing number of recent studies have focused on the differentiation and speciation of insects inhabiting the QTP, e.g., *Gnaptorina* (Li et al., 2021) and *Pseudabris hingstoni* (Wang, 2021). These previous studies have reported associations between speciation patterns and the QTP uplifts (as well as associated climatic changes) and highlight the diversity in phylogeographic structures and divergence processes among taxa. For example, some species used the southeastern QTP as a large refugium during the major glaciations in the Pleistocene (Fan et al., 2012;Liu et al., 2015;Wang et al., 2017), while glacial refugia were maintained in the central platform of the QTP for other native species (Tang et al., 2010;Liu et al., 2013;Muellner-Riehl, 2019).

Grassland caterpillars (Lepidoptera: Erebidae: Lymantriinae: *Gynaephora*) are among the most damaging insect pests to alpine meadows of the QTP (Zhang and Yuan, 2013;Yuan et al., 2015). Eight QTP *Gynaephora* species have been described based on morphological characteristics, accounting for over half of the 15 known *Gynaephora* species worldwide. Therefore, the QTP is recognized as the speciation center of *Gynaephora*, though divergence patterns and speciation mechanisms of the genus in the QTP are largely unknown. Our previous phylogenetic analysis and divergence time estimates for samples from 15 geographic populations based on two mitochondrial genes (*cox1* and *nad5*) and two nuclear genes (*EF-1*α and *GAPDH*) have suggested an important impact of the QTP uplift and associated climatic changes during the late Miocene/early Pleistocene on diversification and speciation. However, species delimitation did not recover all eight *Gynaephora* species based on morphology, i.e., individuals of three *Gynaephora* species showed admixture and were characterized by very low genetic distances (Yuan et al., 2015). It is noteworthy that these three species show high similarity in morphological characteristics (e.g., overall size, wing color, and external genitalia shape) and *G. rouergensis* and *G. minora* are typically sympatric (Zhang and Yuan, 2013). Therefore, sampling of more geographic populations and individuals is necessary to explore species statuses, speciation, and biogeographic patterns of QTP *Gynaephora*.

Here, we conducted extensive sampling from 39 geographic populations, covering the majority of the distribution of each of the eight QTP *Gynaephora* species to investigate phylogeography and speciation based on two mitochondrial genes (*cox1* and *nad5*). We aimed to address the following three questions. (1) How many *Gynaephora* species are in the QTP? (2) Do QTP *Gynaephora* species have high genetic diversity? (3) What are the speciation patterns in QTP *Gynaephora*. This study provides the largest phylogeographic analysis of QTP *Gynaephora* and expands our general understanding of the diversity and speciation of the QTP insects.

## Materials and methods

### Sampling and DNA extraction

A total of 488 individual specimens from 39 geographical localities were used in this study, of which 145 individuals from 15 populations were retrieved from our previous study (Yuan et al., 2015). Our sampling covered most of the distribution of grassland caterpillars in the QTP meadows, with detailed sampling information provided in Figure 1 and Table S1. All sampled specimens were stored in absolute ethyl alcohol in the field and transferred to -20 until they were used for DNA extraction. Total genomic DNA was extracted from each specimen using the DNA Extraction Kit (Tiangen, Beijing, China) following the manufacturer’s protocol.

**Figure 1.**
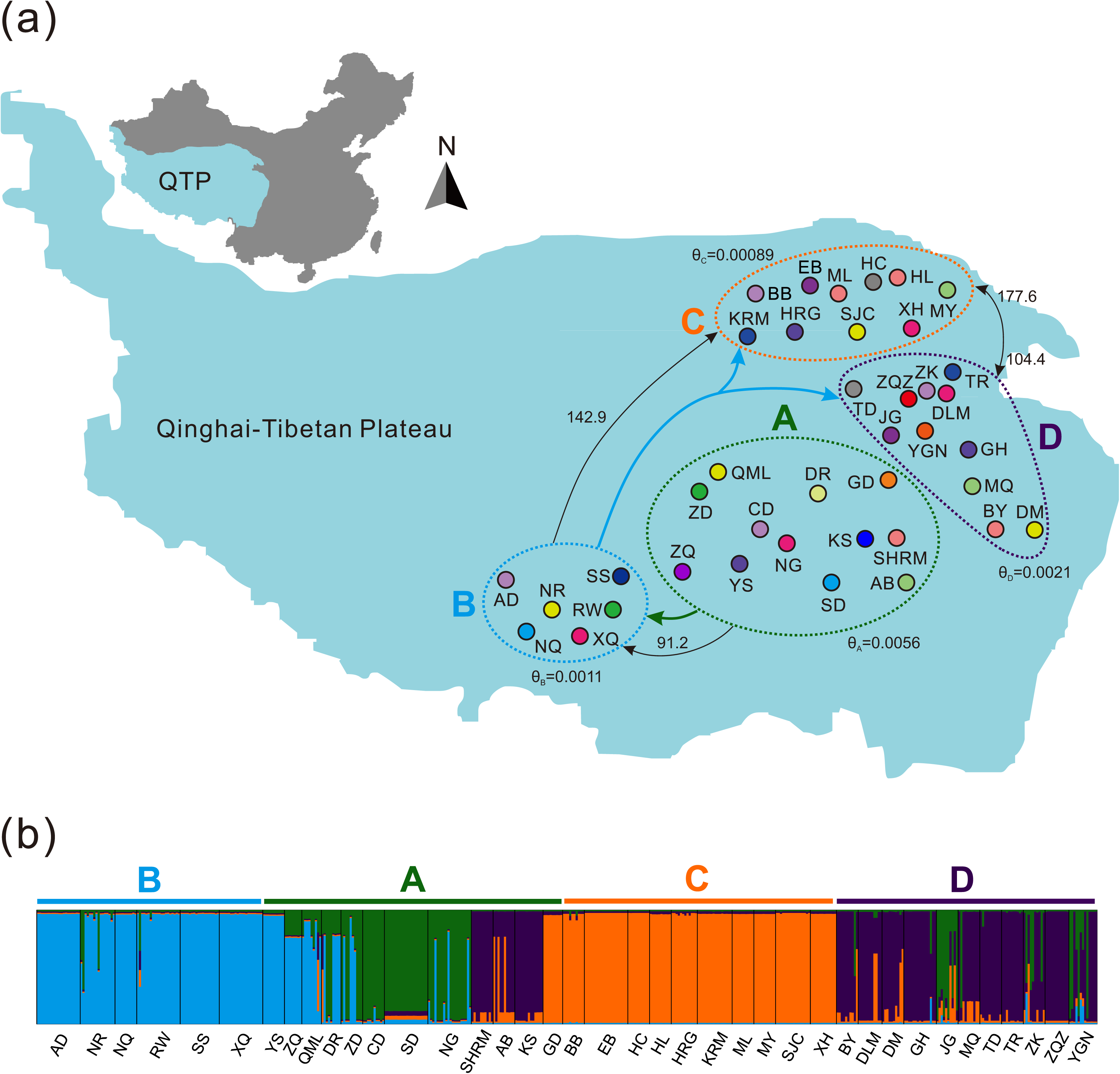
Sampling sites of 39 geographical populations and population structure of the QTP *Gynaephora* species based on the two mitochondrial genes (*cox1* and *nad5*). (a) Sampling sites. Four groups are indicated by A–D, corresponding to four clades in Figures 2 and 3. Bold arrows indicate the dispersal routes inferred by a biogeographic analysis using RASP. Black arrows indicate the direction of gene flow and migration rates are provided above the lines. Effective population sizes (θ) are shown for Clades A–D. (b) Population structure estimated using Structure by setting *K* = 4. Each thin vertical line corresponds to a single population. Four genetic clusters are indicated by A–D, corresponding to four clades in Figures 2 and 3.

### PCR amplification and sequencing

Two mitochondrial genes (*cox1* and *nad5*) were amplified using the same primers and procedures described in our previous study (Yuan et al., 2015). All PCR products were purified using a DNA Gel Purification Kit (Omega, Norwalk, CT, USA) and sequenced in both directions by Sanger sequencing using the same PCR primers.

Sequences of the two mitochondrial genes (*cox1* and *nad5*) were initially aligned using ClustalW in MEGA 5.10 (Tamura et al., 2011) with default parameters to check for stop codons or indels, which could reveal mitochondrial pseudogenes. All sequences newly obtained in this study have been deposited in GenBank under accession numbers OP574353–OP574695 and OP577499–OP577841. Then, sequences of *cox1* and *nad5* were concatenated using DAMBE 7.2.6 (Xia, 2018). Mitochondrial haplotypes were identified using DNASP 5.10 (Librado and Rozas, 2009).

### Phylogenetic analysis and population structure

For each gene, substitution saturation was evaluated using DAMBE 7.2.6 (Xia, 2018). A lack of evidence for saturation indicated that all codon positions can be used for the phylogenetic analysis. The best partitioning schemes and corresponding nucleotide substitution models for the dataset (*cox1* + *nad5*) was selected by PartitionFinder 1.1.1 (Lanfear et al., 2012) and used for phylogenetic analyses, as in our previous study (Yuan et al., 2015). The phylogenetic tree was constructed based on the mitochondrial dataset (*cox1*+*nad5*) using maximum likelihood (ML) and Bayesian inference (BI). Both BI and ML analyses were conducted using the CIPRES Science Gateway 3.3 (Miller et al., 2010). The ML analysis was carried out using RAxML-HPC2 on XSEDE 8.0.24 (Stamatakis, 2014) with the GTRGAMMA model, and 1,000 bootstrap (BS) replicates were used to estimate node reliability. The BI analysis was conducted using MrBayes 3.2.2 (Ronquist et al., 2012) on XSEDE. Four Markov chains (three hot and one cold chain) were independently run two times for 1×10^6^ generations, with sampling every 1,000 generations. Chain congruence was assessed by the effective sample size (ESS) (>100) and the potential scale reduction factor (PSRF) (approximately 1.0), as recommended in MrBayes 3.2.2 documentation (Ronquist et al., 2012). The first 25% of samples were discarded as burn-in, and the remaining samples were used to construct a 50% majority-rule consensus tree. Bayesian posterior probabilities (PP) were calculated.

A median-joining network was generated for all mitochondrial haplotypes using Network 4.6.0.0 (Bandelt et al., 1999). Population structure was further assessed by a Bayesian model-based method implemented in Structure 2.3 (Pritchard et al., 2000;Falush et al., 2003). The admixture model in Structure was used and the number of clusters (*K*) was varied from 2 to 8. For each value of *K* (putative number of populations), 300,000 Markov chain Monte Carlo (MCMC) iterations were run, following a burn-in of 100,000 iterations. Structure Harvester (Earl and vonHoldt, 2012) was used to determine the optimum value of clusters (*K*) by a comparison of mean log probability LnP(*K*) and by calculating the Δ*K* value for each *K* value (EVANNO et al., 2005).

### Genetic diversity and differentiation

Haplotype diversity (*h*) and nucleotide diversity (π) for all 39 geographic populations and each of the four clades (based on the population structure analysis) were calculated using DnaSP 5.10 (Librado and Rozas, 2009). Genetic differentiation (*F*_ST_) between the 39 geographic populations was calculated using Arlequin 3.5 (Excoffier and Lischer, 2010). The significance of *F*_ST_ between populations was tested by 10,000 permutations in Arlequin 3.5 (Excoffier and Lischer, 2010). To investigate genetic structuring between sampling locations, an analysis of molecular variance (AMOVA) (Excoffier et al., 1992) was performed with Arlequin 3.5 (Excoffier and Lischer, 2010). Mantel tests were performed using Isolation By Distance Web Service (IBDWS) (Jensen et al., 2005) to test the correlation between genetic distance and geographic distance.

### Population demography

The historical population demographics of QTP *Gynaephora* species was investigated by neutrality tests (Fu’s *Fs*, Fu and Li’s *F*\* and *D*\*, and Tajima’s D) and pairwise mismatch distributions for all 39 sampling locations combined and each of four clades separately using DnaSP 5.10 (Librado and Rozas, 2009). Fu’s *Fs*, Fu and Li’s *F*\* and *D*\*, and Tajima’s *D* were calculated to detected population demography (Tajima, 1989;Fu, 1997). Significant negative *F*_S_ values can be taken as evidence for expansions, while positive values might result from population subdivision or a recent population bottleneck (Fu, 1997). For the mismatch distribution analysis, a unimodal distribution indicates a recent demographic expansion, while multimodal distributions indicate population stability (Rogers and Harpending, 1992;Harpending et al., 1998).

### Divergence time estimation

Divergence times of the QTP *Gynaephora* were estimated by a molecular clock approach implemented in BEAST 1.8.1 (Drummond et al., 2012) using the phylogenetic tree of haplotypes as a constraint tree. A lognormal prior was used for *cox1* and the mean substitution rate was set to 0.0115 per site per million years (Ma), while the substitution rate for *nad5* was scaled to the mean *cox1* rate (according to the K2P genetic distances between *cox1* and *nad5*), following the method described in our previous study (Yuan et al., 2018). The following settings were used: linked tree model, HKY+I substitution model for each mitochondrial gene, strict clock, Yule model of speciation, and default values for the remaining parameters. Chain convergence was checked using the ESS implemented in Tracer 1.5. Posterior estimates of *Gynaephora* ages and 95% highest posterior densities (HPD) were summarized using TreeAnnotator v1.8.1 (Drummond et al., 2012). All BEAST analyses were conducted on the CIPRES Science Gateway 3.3 (Miller et al., 2010).

### Biogeographical analysis

Ancestral area reconstruction was conducted to investigate the biogeographical history of *Gynaephora* by using a Bayesian binary MCMC (BBM) method, as implemented in RASP 3.1 (Yu et al., 2015). Seven biogeographical areas were defined based on the species distributions and results of the phylogenetic analysis (Figure 1): the distributions of (a) *G. groenlandica*; (b) *G. rossii*; (c) *G. selenitica*; and (d–g) the four clades A–D in Figure 1. The BBM analysis was run with the following settings: 10001 credible trees generating from the BEAST analysis were used, the maximum number of areas for all nodes was set to seven, and the remaining parameters were set to default values.

### Modelling gene flow

Gene flow between the four clades (A–D in Figure 1) identified by phylogenetic analyses was modeled by using a coalescent-based approach implemented in Migrate-n (Beerli, 2006;Beerli and Palczewski, 2010). This software estimates long-term average values of gene flow and can be used to test predictions about refugial structure and the direction of migration from putative refugia. The Sanjiangyuan region, locating in the central and eastern QTP, might represent a refugium of the QTP *Gynaephora* during glacial periods, as has been inferred in previous studies of other taxa (Yang et al., 2009;Qiu et al., 2011). Therefore, to reduce the number of models, we first divided the 39 populations into two groups based on the phylogenetic results, i.e., group A (Clade A in Figure 1) and group BCD (Clades B–D in Figure 1), and constructed three migration models (M1–M3 Figure S2). Then, we tested 14 migration models for three groups (Clade A, Clade B, and Clades C+D) and 9 migration models for four groups (Clades A–D in Figure 1, Figure S2).

Migrate-n was used to estimate migration rates (M) and effective population size (θ) parameters as well as the marginal likelihood of each gene flow model (Beerli, 2006). The gene flow models were ranked according to log Bayes factors (LBF) calculated from the Bezier corrected marginal likelihoods of the data given the model. The marginal likelihoods for each model were approximated by thermodynamic integration of the MCMC over four heated chains (Metropolis coupled MCMC), as described previously (Beerli and Palczewski, 2010). We used a uniform prior for θ between 0 and 0.1 and a sampling window of 0.01 on which to generate new proposals; for M, we used a uniform prior between 0 and 1000 with a window of 100 model. A four-chain static heating scheme was used, and 100,000 trees were discarded as burn-in before recording 1000,000 steps with an increment of 100, resulting in 40 million samples. To evaluate convergence, we evaluated ESS (>10000) and the posterior distributions of the parameters to determine whether they were unimodal smooth curves. Bayes factors (BF) were calculated as a ratio of the marginal likelihoods to calculate model probabilities.

## Results

### Population structure

A total of 40 haplotypes were detected for the combined mitochondrial gene dataset (*cox1* and *nad5*) (Table S1). Among these haplotypes, 29 haplotypes were specific to a single population, and the remaining 11 haplotypes were shared by multiple geographic populations (Table S1, Figure 2). Haplotypes H3 and H4 showed the highest frequencies, followed by H2, H5, H7, and H11 (Figure 2). Only one haplotype was detected in all eighteen populations, and the DM population (six haplotypes) had the largest number of haplotypes (Table S1).

**Figure 2.**
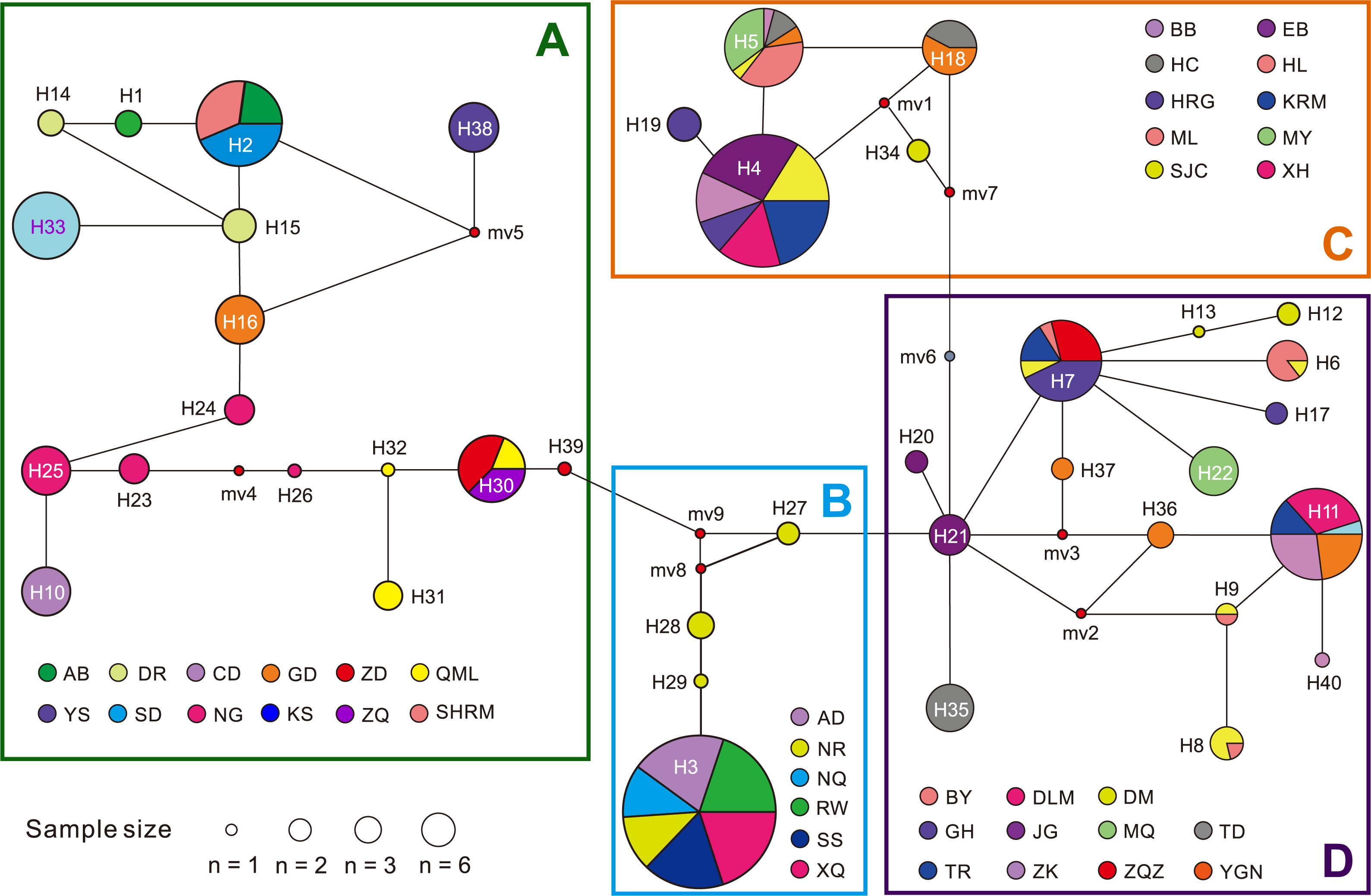
Median-joining network of 40 haplotypes of QTP *Gynaephora* species. Four groups are indicated by A–D, corresponding to four clades in Figures 1 and 3. See Table S1 for full names of the 39 geographical populations. The size of each circle (pie chart) indicates the number of individuals sharing this haplotype. The nine smallest circles (mv1–mv9) indicate hypothesized haplotypes.

Phylogenetic analyses based on 40 haplotypes using two methods (BI and ML) yielded nearly identical topologies (Figure 3). The QTP *Gynaephora* species formed a strongly supported monophyletic group (PP = 1.0, BS = 100) and consisted of four clades (A–D in Figures 1 and 3). Clade A (16 haplotypes in 12 populations) was in the basal position, and clade B (4 haplotypes in 6 populations) was sister to clade C (5 haplotypes in 10 populations) and clade D (15 haplotypes in 11 populations) (Figure 3). These four clades were also obtained by a haplotype network analysis (Figure 2).

**Figure 3.**
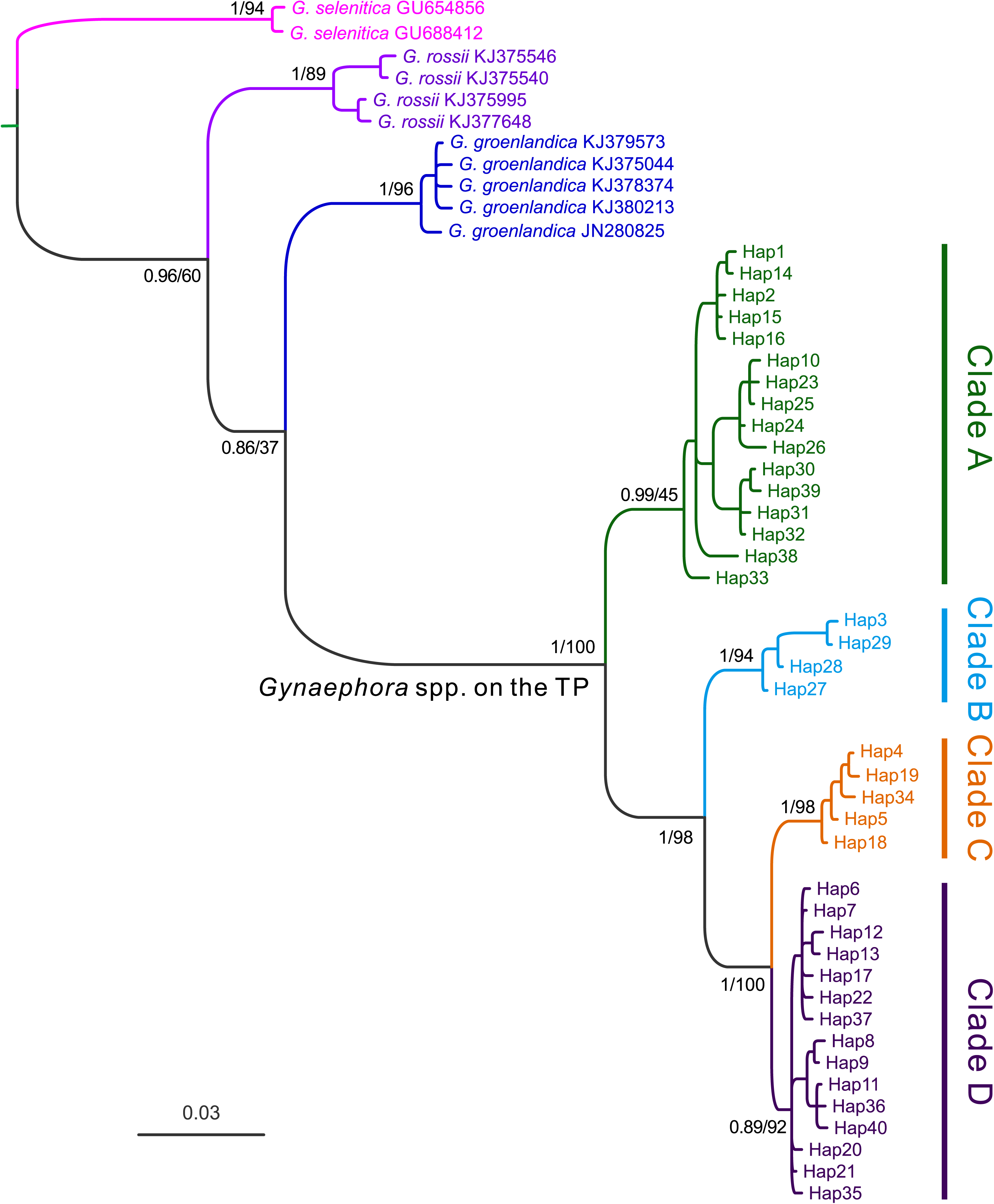
Bayesian phylogenetic trees of *Gynaephora* species based on two mitochondrial genes (*cox1* and *nad5*). Posterior probabilities (left) and bootstrap support (right) values are shown above the main nodes.

In a structure analysis, the assignment of individuals to four groups was best supported, with a maximum log likelihood of posterior probability (Ln P(*K*) =-2283.22) and a maximum Δ*K* (Δ*K* = 38.55) at *K* = 4 (Figure 1b). These four groups (A–D) were largely congruent with those obtained in phylogenetic and network analyses (Figures 2 and 3). Individuals from groups B and C were well assigned to clusters with a small number of admixed individuals, while high levels of genetic admixture were found in groups A and D (Figure 1b).

### Genetic diversity and differentiation

For QTP *Gynaephora* species overall, high levels of haplotype diversity (*h* = 0.913 ± 0.007) and nucleotide diversity (π = 2.288 ± 0.033) were observed (Table 1). Among the four clades (Figures 1–3), Clade A showed the highest genetic diversity (*H* =16, *h* = 0.888 ± 0.012, π = 0.641 ± 0.019%), followed by Clade D (*H* =15, *h* = 0.837 ± 0.020, π = 0.201 ± 0.006%) and Clade C (*H* = 5, *h* = 0.567 ± 0.040, π = 0.061 ± 0.006%), while Clade B (*H* = 4, *h* = 0.112 ± 0.042, π = 0.073 ± 0.30%) showed the lowest genetic diversity (Table 1).

**Table 1.** Genetic diversity and neutrality tests for 39 geographic populations combined and for each of four clades of QTP *Gynaephora*. *n*, sample size sequenced. *H*, number of haplotypes. *h,* haplotype diversity. π, nucleotide diversity. * *P* < 0.05, ** *P* < 0.001, ^ns^ *P* > 0.05. See Figure 1 for the four clades (Clades A–D).

| | $n$ | $H$ | $h$ | $\pi$ (%) | Tajima's $D$ | Fu and Li's $D^*$ | Fu and Li's $F^*$ | Fu's $F_S$ |
| --- | --- | --- | --- | --- | --- | --- | --- | --- |
| All populations | 488 | 40 | $0.913 \pm 0.007$ | $2.288 \pm 0.033$ | $2.98^*$ | $2.73^*$ | $3.42^*$ | $24.05^{**}$ |
| Clade A | 138 | 16 | $0.888 \pm 0.012$ | $0.641 \pm 0.019$ | $1.31^{\text{ns}}$ | $1.64^*$ | $1.81^*$ | $4.36^*$ |
| Clade B | 104 | 4 | $0.112 \pm 0.042$ | $0.073 \pm 0.030$ | $-1.45^{\text{ns}}$ | $1.42^{\text{ns}}$ | $0.47^{\text{ns}}$ | $1.25^{\text{ns}}$ |
| Clade C | 126 | 5 | $0.567 \pm 0.040$ | $0.061 \pm 0.006$ | $0.10^{\text{ns}}$ | $0.92^{\text{ns}}$ | $0.77^{\text{ns}}$ | $-0.06^{\text{ns}}$ |
| Clade D | 120 | 15 | $0.837 \pm 0.020$ | $0.201 \pm 0.006$ | $0.62^{\text{ns}}$ | $0.73^{\text{ns}}$ | $0.82^{\text{ns}}$ | $-2.57^*$ |

Almost all pairwise *F*_ST_ values between the 39 sampling sites were significantly greater than 0 (Table S2), indicating substantial genetic differentiation between QTP *Gynaephora* populations. Mantel tests of the 39 QTP *Gynaephora* populations indicated that there is a significant positive correlation between genetic distances and geographic distances (*r* = 0.169–0.517, *P* < 0.01; Table S3). Among the four clades, Mantel tests also revealed a significant correlation between genetic and geographic distances for Clade A (*r* = 0.191–0.457, *P* < 0.05; Table S3).

To further investigate genetic differentiation and genetic structuring between the 39 geographic populations we performed AMOVA of the four clades according to phylogenetic and network analyses (Figures 1 and 2). A significantly positive *F*_CT_ (*F*_CT_ = 0.8701, *P* < 0.001; Table 2) suggested that geographic structuring existed among the examined populations. Genetic differentiation among the four clades accounted for 87.01% of the total genetic variance, whereas differentiation within clades only accounted for 1.35% of genetic variation, indicating low gene flow among the four clades. We also performed an additional AMOVA by pooling the 39 geographic populations into 2–7 groups according to altitude of sampling sites (Table 2). The significant positive *F*_CT_ (*F*_CT_ = 0.3882–0.5431, *P* < 0.001; Table 2) and high percentage of variance explained (98.46–98.71% among and within groups) revealed the impact of altitude on the population genetic structure of the QTP *Gynaephora* species.

**Table 2.** Results of analysis of molecular variance (AMOVA). ^a^, grouped according to the results of phylogenetic and network analyses (Figures 1 and2); ^b^, grouped by altitude. * *P* < 0.001.

| Group | Source of variation | df | Sum of squares | Variance components | Percentage of variation (%) | *F*-statistic |
| --- | --- | --- | --- | --- | --- | --- |
| Four groups<sup>a</sup> | Among groups | 3 | 6019.55 | 16.23 | 87.01 | $F_{CT} = 0.8701^*$ |
| | Among populations within groups | 35 | 947.68 | 2.17 | 11.64 | $F_{SC} = 0.8960^*$ |
| | Within populations | 449 | 113.08 | 0.25 | 1.35 | $F_{ST} = 0.9865^*$ |
| Two groups<sup>b</sup> | Among groups | 1 | 2274.71 | 9.04 | 46.43 | $F_{CT} = 0.4643^*$ |
| | Among populations within groups | 27 | 4692.51 | 10.17 | 52.27 | $F_{SC} = 0.9758^*$ |
| | Within populations | 449 | 113.08 | 0.25 | 1.29 | $F_{ST} = 0.9870^*$ |
| Three groups<sup>b</sup> | Among groups | 2 | 2432.04 | 6.67 | 39.10 | $F_{CT} = 0.3909^*$ |
| | Among populations within groups | 36 | 4535.18 | 10.14 | 59.43 | $F_{SC} = 0.9757^*$ |
| | Within populations | 449 | 113.08 | 0.25 | 1.48 | $F_{ST} = 0.9852^*$ |
| Four groups<sup>b</sup> | Among groups | 3 | 3376.96 | 9.07 | 51.63 | $F_{CT} = 0.5162^*$ |
| | Among populations within groups | 35 | 3590.27 | 8.24 | 46.94 | $F_{SC} = 0.9703^*$ |
| | Within populations | 449 | 113.08 | 0.25 | 1.43 | $F_{ST} = 0.9856^*$ |
| Five groups<sup>b</sup> | Among groups | 4 | 2860.53 | 6.34 | 38.83 | $F_{CT} = 0.3882^*$ |
| | Among populations within groups | 34 | 4106.70 | 9.74 | 59.63 | $F_{SC} = 0.9748^*$ |
| | Within populations | 349 | 113.08 | 0.25 | 1.54 | $F_{ST} = 0.9845^*$ |
| Six groups<sup>b</sup> | Among groups | 5 | 3979.05 | 9.03 | 54.31 | $F_{CT} = 0.5431^*$ |
| | Among populations within groups | 33 | 2988.18 | 7.35 | 44.18 | $F_{SC} = 0.9669^*$ |
| | Within populations | 449 | 113.08 | 0.25 | 1.51 | $F_{ST} = 0.9849^*$ |
| Seven groups<sup>b</sup> | Among groups | 6 | 4046.13 | 8.81 | 53.58 | $F_{CT} = 0.5358^*$ |
| | Among populations within groups | 32 | 2921.10 | 7.38 | 44.88 | $F_{SC} = 0.9670^*$ |
| | Within populations | 449 | 113.08 | 0.25 | 1.53 | $F_{ST} = 0.9846^*$ |

### Population demography

The mismatch distribution for the 40 haplotypes of the 39 geographic populations was multimodal (Figure S1), indicating that no obvious expansion occurred in the QTP *Gynaephora* species. Similar results were obtained for three clades (Clades A, B, and D), whereas Clade C showed a unimodal distribution, which could be due to either demographic expansion (population expansion) or to positive selection. In neutrality tests, significant negative values were not obtained for QTP *Gynaephora* as a whole or for Clades A, B, and C (Table 1). Significant positive values for each of four neutrality tests were found in all populations combined, whereas Fu’s *F*_S_ value was significantly negative in Clade D (Table 1).

### Divergence time estimation

Divergence time estimates for the genus *Gynaephora* based on a strict mitochondrial molecular clock are shown in Figure 4. Divergence within the genus was dated to the late Miocene, approximately 6.05 Ma (95% HPD: 2.9–10.83 Ma). The divergence of QTP *Gynaephora* species from non-QTP species likely occurred during the early Pliocene, at around 4.65 Ma (95% HPD: 2.22–8.12 Ma). The earliest split within QTP *Gynaephora* was dated to the early Pleistocene at about 2.08 Ma (95% HPD: 0.90–3.68 Ma), followed by further splitting at around 1.31 Ma (95% HPD: 0.58–2.39 Ma). Clades C and D diverged about 0.53 Ma (95% HPD: 0.2–1.03 Ma).

**Figure 4.**
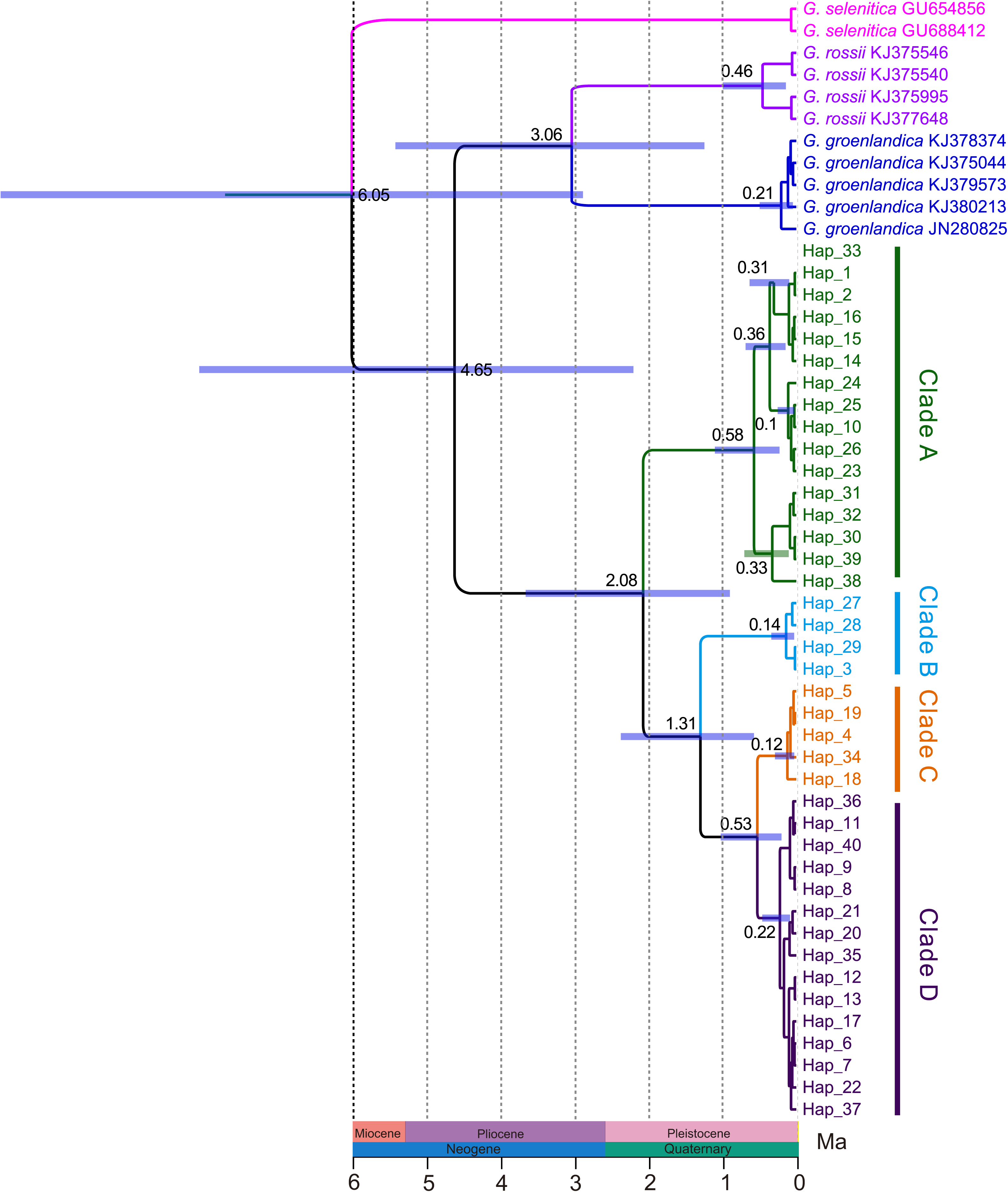
Divergence time estimation of the *Gynaephora* species using BEAST. Blue bars represent the 95% highest probability density interval. The numbers above nodes are the mean divergence times. Ma, million years ago.

### Biogeographic reconstruction

The BBM biogeographic analysis revealed that the genus *Gynaephora* likely originated in central Europe (see Figure 5c for the distribution of the type species *G. selenitica*) with high probability; 12 dispersal and 6 vicariance events led to the colonization of current ranges with two expansion routes: northward dispersal to form several arctic *Gynaephora* species (e.g., *G. rossii* and *G. groenlandica*) and southward dispersal to form the QTP *Gynaephora* species (Figure 5). The common ancestor of the QTP *Gynaephora* species likely first dispersed to the Sanjiangyuan Region of the QTP (Clade A in Figure 1) and then to the Naqu Region, leading to the emergence of *G. alpherakii* by two westward dispersal events and one vicariance event, followed by dispersal to the North and Northeast QTP (Figure 5) and the split into two clades (C and D in Figure 1).

**Figure 5.** Biogeographic analysis of *Gynaephora* species using RASP. Clades A–D are the same as those in Figures 1–3. Present-day ranges for each haplotype are color-coded and drawn as small circles at the ends of terminal branches before haplotype names. Pie charts at internal nodes represent marginal probabilities for each alternative ancestral range (color codes).

### Gene flow

A total of 26 models were evaluated using Migrate-n (Table 3, Figure S2). The best-supported models were M3 for two groups (A, BCD), M5 for three groups (A, B, CD), and M20 for four groups (Clades A–D) (Table 3, Figure S2). These best models consistently supported Clade A as the refugium with unidirectional gene flow from Clade A to Clade B (mean *Nm* = 91.2) and from Clade B to Clade C (mean *Nm* = 142.9) (Figure 1, Table S4). Bidirectional and asymmetrical gene flow were found between Clades C and D (Figure 1, Table S4). Theta estimates suggested that *Ne* was highest in Clade A and lowest in Clade C (Table S4).

**Table 3.** Results from model comparisons tested in Migrate-n. Model descriptions are provided in Figure S2. The number of parameters for each model, Bezier approximation scores of log marginal likelihoods (l mL), log Bayes factors (LBF), and model probabilities are shown. A–D correspond to Clades A–D in Figure 1.

| Model description | Model | Parameters | Bezier l mL | LBF | Probability |
| --- | --- | --- | --- | --- | --- |
| Two groups (A, BCD) |  |  |  |  |  |
| Full model | M1 | 4 (2θ, 2M) | -5802.97 | 340.82 | <0.01 |
| BCD as refugium, 1 route | M2 | 3 (2θ, 1M) | -5646.05 | 26.98 | <0.01 |
| A as refugium, 1 route | M3 | 3 (2θ, 1M) | -5632.56 | 0 | 1 |
| Three groups (A, B, CD) |  |  |  |  |  |
| Full Model | M4 | 9 (3θ, 6M) | -4968.95 | 456.0 | <0.01 |
| A as refugium, 1 route | M5 | 5 (3θ, 2M) | -4740.95 | 0 | 1 |
| A as refugium, 2 routes | M6 | 6 (3θ, 3M) | -4811.54 | 141.18 | <0.01 |
| A as refugium, 2 routes | M7 | 6 (3θ, 3M) | -4872.75 | 263.6 | <0.01 |
| A as refugium, 2 routes | M8 | 7 (3θ, 4M) | -4838.25 | 194.6 | <0.01 |
| A as refugium, 1 route | M9 | 5 (3θ, 2M) | -4785.92 | 89.94 | <0.01 |
| A as refugium, 2 routes | M10 | 6 (3θ, 3M) | -4860.63 | 239.36 | <0.01 |
| A as refugium, 4 routes | M11 | 8 (3θ, 5M) | -4842.57 | 203.24 | <0.01 |
| A as refugium, 3 routes | M12 | 7 (3θ, 4M) | -4854.30 | 226.7 | <0.01 |
| A as refugium, 2 routes | M13 | 5 (3θ, 2M) | -4816.21 | 150.52 | <0.01 |
| A as refugium, 2 routes | M14 | 6 (3θ, 3M) | -4864.94 | 247.98 | <0.01 |
| A as refugium, 2 routes | M15 | 6 (3θ, 3M) | -4810.94 | 139.98 | <0.01 |
| A as refugium, 2 routes | M16 | 7 (3θ, 4M) | -4858.42 | 234.94 | <0.01 |
| A as refugium, 2 routes | M17 | 6 (3θ, 3M) | -4838.33 | 194.76 | <0.01 |
| Four groups (A, B, C, D) |  |  |  |  |  |
| Full model, 4 populations | M18 | 16 (4θ, 12M) | -5463.89 | 1464.78 | <0.01 |
| A as refugium, 1 route | M19 | 7 (4θ, 3M) | -4744.14 | 25.28 | <0.01 |
| A as refugium, 2 routes | M20 | 8 (4θ, 4M) | -4731.50 | 0 | 1 |
| A as refugium, 1 route | M21 | 7 (4θ, 3M) | -4782.24 | 101.48 | <0.01 |
| A as refugium, 2 routes | M22 | 8 (4θ, 4M) | -4842.89 | 222.78 | <0.01 |
| A as refugium, 2 routes | M23 | 7 (4θ, 3M) | -4800.06 | 137.12 | <0.01 |
| A as refugium, 3 routes | M24 | 8 (4θ, 4M) | -4864.18 | 265.36 | <0.01 |
| A as refugium, 2 routes | M25 | 8 (4θ, 4M) | -4933.58 | 404.16 | <0.01 |
| A as refugium, 3 routes | M26 | 9 (4θ, 5M) | -4805.27 | 147.54 | <0.01 |

## Discussion

### The number of QTP *Gynaephora* species may be overestimated

The unique geographical and environmental conditions of the QTP led to a large number of endemic species. *Gynaephora* is a small genus, with only one species described in China before the 1970s and eight species described to date according to morphological characteristics (Zhang and Yuan, 2013). However, these eight morphological species were not well-supported by molecular data in our previous study (Yuan et al., 2015). In this study, by extensive sampling covering almost the whole geographical distribution and phylogenetic analyses, the number of valid species was further reduced to four monophyletic groups (Clades A–D in Figure 1). Clades B and C corresponded to *G. alpherakii* and *G. menyuanensis*, respectively. Clades A and D each contained multiple morphologically described species. Among three species included in Clade A, *G. qinghaiensis* had a wide geographical distribution, while the two other *Gynaephora* (*G. qumalaiensis* and *G. jiuzhiensis*) are locally distributed (Yuan et al., 2015). *G. qinghaiensis* has long been considered the only species in alpine meadows of the QTP, i.e., grassland caterpillars destroying alpine meadows are all considered *G. qinghaiensis*. Although the Latin name *G. alpherakii* is frequently used to describe the QTP grassland caterpillars in the literature, Chou et al. (Chou and Ying, 1979) found that no specimens of grassland caterpillars sampled conformed to the morphological characteristics of *G. alpherakii*, questioning the existence of this species in the QTP. However, both our results and previous results consistently indicated that grassland caterpillars collected from multiple geographical localities in the Naqu Region of Tibet are an independent species, including those in Amdo County, the locality of the type specimen of *G. alpherakii*. Further studies including morphological characteristics are necessary to confirm whether this recovered monophyletic group is indeed *G. alpherakii,* as originally described.

Three *Gynaephora* species (*G. aureata*, *G. rouergensis,* and *G. minora*) clustered together to form Clade D, as found in our previous study (Yuan et al., 2015). *G. aureata* is widely distributed, while the two other species are only distributed in Ruoergai County (Chou and Ying, 1979;Zhang and Yuan, 2013). The two sympatric species are morphologically similar, with obvious differences only in wing size (Chou and Ying, 1979), likely due to phenotypic plasticity. In addition, a sympatric distribution may facilitate relatively strong mitochondrial introgression due to interspecific hybridization. Among eight QTP *Gynaephora* species, *G. menyuanensis* is the most recently described and is mainly distributed in the northeastern QTP (Zhang and Yuan, 2013). This species can be effectively differentiated based on several morphological characteristics from other *Gynaephora* species. The validity of this species was also supported by molecular data in our present and previous study, suggesting the independent species status of *G. menyuanensis*.

### High genetic diversity in QTP *Gynaephora* species

Genetic diversity is a fundamental component of biodiversity. The higher the genetic diversity within a species, the greater the potential for adaptation in response to changing environments. Generally, genetic diversity can be measured by two indices, i.e., haplotype diversity (h) and nucleotide diversity (π). Nucleotide diversity represents the cumulative degree of genetic variation during evolution, while haplotype diversity reflects the probability that two randomly sampled alleles are different and is more relevant to population dynamics at short time scales (Pauls et al., 2013;Leitwein et al., 2020). Patterns of genetic diversity can be assigned to four categories (Grant and Bowen, 1998): (1) low *h* and low π, (2) high *h* and low π, (3) low *h* and high π, and (4) high *h* and high π. The results for all 39 *Gynaephora* populations as a whole were consistent with the fourth category, i.e., high *h* (>0.5) and high π (>0.5%), suggesting that QTP *Gynaephora* was a large stable population with a long evolutionary history or secondary contact between differentiated lineages. High level of genetic diversity may be necessary for QTP *Gynaephora* species to adapt to the extreme high-altitude environments. This also provides a genetic basis for maintaining a high population density and explains frequent outbreaks of grassland caterpillars in alpine meadows of the QTP.

Among the four clades, Clade A showed the highest haplotype and nucleotide diversities, inconsistent with the hypothesis that source populations generally have low genetic diversity due to the founder effect. Given that Clade A consisted of three morphologically described species, we proposed that the high level of genetic diversity found in Clade A may be related to secondary contact between these differentiated species/populations during glacial-interglacial cycle periods. Clade B (*G. alpherakii*) fell into the first category (low *h* and low π), suggesting a contribution of founder events in which new populations were established by a small number of individuals drawn from the large ancestral Clade A. In particular, only one exclusive haplotype was found in geographical populations AD, NQ, RW, SS, and XQ from Clade B, further confirming that genetic drift might eliminate many haplotypes, leaving only one haplotype in these populations. It is possible that clade B, found in higher-altitude areas, faced stronger environmental pressures, resulting in lineage-specific haplotypes. Since 1960s, various insecticides have been used extensively and frequently for controlling *Gynaephora* (Zhang and Yuan, 2013), which may result in reduced levels of genetic variation in some populations. Therefore, population bottlenecks resulting from widespread insecticide application likely contributed to the low genetic diversity of Clade B.

Both Clades C and D were consistent with the second category, i.e., high haplotype diversity (*h* > 0.5) and low nucleotide diversity (π < 0.5%). This might be attributed to population expansion after population bottleneck followed by rapid population growth and he accumulation of new mutations (Grant and Bowen, 1998). High haplotype diversity was likely related to the high mitochondrial DNA mutation rate, as indicated by a haplotype network analysis (Figure 2) with 1–2 mutation steps before the generation of a new haplotype. This high mutation rate may be also associated with the extensive use of insecticides and may result in an increased rate of development of resistance genes, making effective control challenging. Further explorations are needed to clarify differences in genetic diversity among the four clades of the QTP *Gynaephora* species.

### Isolation and subsequent divergence is the key speciation mechanism of the QTP *Gynaephora* species

The divergence between QTP *Gynaephora* species and non-QTP species was estimated to occur in the early Pliocene (around 4.65 Ma), which was slightly younger our previous estimate (Yuan et al., 2015). The genus *Gynaephora* likely originated in central Europe, as indicated by the biogeographic analysis. Therefore, ancestors of QTP *Gynaephora* arrived at the QTP during a period of uplift. The QTP has experienced uplifts many times; an intensive uplift probably began during the Miocene and the most intensive uplifts during the Pliocene and Pleistocene are known as the Qingzang Movement (3.6–1.7 Ma) and the Kun-Huang Movement (1.2–0.6 Ma) (Shi et al., 1998;Zheng et al., 2000;An et al., 2001). These intensive uplifts of the QTP were likely causes of interspecific divergence in QTP *Gynaephora* date to the early Pleistocene to Pliocene (2.08–0.53Ma). In addition, changes in vegetation from forests to grasslands due to climate change during the Pleistocene (Wu et al., 2001) not only provided abundant food resources, an important basis for population expansion, but also likely provided new habitats, contributing to population differentiation and speciation. Therefore, we proposed that grassland caterpillars arrived at the QTP after the QTP uplifts and then gradually diverged, resulting in speciation influenced by plateau uplifts and associated climatic fluctuations.

Biogeographic analysis, divergence time estimation, and gene flow analysis showed that the region where *Gynaephora* first arrived was the interior of the QTP (Sanjiangyuan region, Clade A in Figure 1) after its migration from Eurasia. This was also supported by the higher genetic diversity in Clade A than in the three other clades. Given the unidirectional gene flow from Clade A to Clade B, the QTP *Gynaephora* may have remained in the interior of the QTP during the glacial period, rather than dispersing to low-altitude environments. Although it is not clear whether the QTP formed a large ice sheet during the glacial period, some animals did remain in the QTP interior during these periods. Therefore, the Sanjiangyuan region was likely a refuge for QTP *Gynaephora* during glacial periods of the Pleistocene. Generally, postglacial differentiation could induce a decrease in genetic diversity along the expansion route, with an increase in the distance from refugia (Comes and Kadereit, 1998;Provan and Bennett, 2008). This hypothesis can explain the decrease in genetic diversity from Clade A to Clade B; however, it was difficult to explain the increasing trend from Clade B to Clade C. Both Clades C and D inhabited relatively low altitudes and showed relatively high genetic diversity, likely due to a high level of bidirectional gene flow between these two clades during the interglacial period. Significant genetic differentiation among the four clades and unidirectional gene flow (Clade A to Clade B and then Clade C), together with the observed association between genetic and geographic distances, indicated that geographic isolation and subsequent divergence was the dominant mode of speciation in the QTP *Gynaephora*, as has been proposed in some animal taxa endemic to the QTP (Lei et al., 2014;Wen et al., 2014;Favre et al., 2015).

## Conclusion

In this study, we investigated the phylogeography and speciation of QTP *Gynaephora* based on two mitochondrial genes by extensive sampling. We recovered four monophyletic clades, indicating that the number of QTP *Gynaephora* species may be overestimated. The taxonomic status of the eight QTP *Gynaephora* species described based on morphological characteristics needs to be further studied by using a combination of morphological and nuclear data. High levels of genetic diversity detected in QTP *Gynaephora* may explain the high-altitude adaptation and outbreaks in alpine meadows of the QTP. The ancestor of the QTP arrived in the region during the Pliocene, when the QTP had been uplifted, and gradually diverged, influenced by intensive plateau uplifts and associated climate fluctuations during the Pliocene and Pleistocene periods. Accordingly, isolation and subsequent divergence may explain speciation in QTP *Gynaephora*. This study is the largest phylogeographic analysis of QTP *Gynaephora* to date and provides insights that may contribute to the control of these pests in alpine meadows.

## Acknowledgments

This study was funded by the National Natural Science Foundation of China (31201520), the Second Tibetan Plateau Scientific Expedition and Research (STEP) Program (2019QZKK0302) and National Science & Technology Fundamental Resources Investigation Program of China (2019FY100400/2019FY100404).

## Author contributions

M.L.Y. conceived and designed the experiments. M.L.Y., M.H.B, Q.L.Z., Z.L.G. and J.W. sampled insect specimens. M.H.B, Q.L.Z. and J.W. conducted experiments. M.L.Y., M.H.B., Q.L.Z., Z.L.G., and M.L. performed data analyses. M.L.Y., M.H.B. and Q.L.Z. wrote the manuscript. All authors read and approved the final version of the manuscript.

## Supplemental material

Supplemental material for this article is available online.

## Supporting Information

**Figure S1** Mismatch distributions for all 39 geographical populations combined and each of the four clades of the QTP *Gynaephora* species. See Figures 1–3 for Clades A–D.

**Figure S2** All the 26 migration models (M1–M26) tested in Migrate-n. The three best models are highlighted by a light orange box. See Figures 1–3 for Clades A–D.

**Table S1** Sampling information and haplotype distributions for 39 geographic populations of QTP *Gynaephora* species.

**Table S2** Pairwise *F*_ST_ (below diagonal) and *P*-values (above diagonal) for the 39 geographic populations of the QTP *Gynaephora* species based on *cox1* and *nad5* data. See Table S1 for population abbreviations.

**Table S3** Mantel tests for relationships between genetic and geographic distances of the QTP *Gynaephora* species. See Figure 1 for Clade A, Clade B, Clade C, and Clade D.

**Table S4** Parameter estimates of Theta (θ) and immigration rate (M) for migration model comparisons based on two mitochondrial genes (*cox1* and *nad5*). Mean values and 2.5–97.5% intervals in brackets are provided for each parameter. Detailed information for the three best models (M3, M5, and M20) is provided in Table 3 and Figure S2.

