## Supplemental Figure S2 for "Mitochondrial phylogeography of grassland caterpillars (Lepidoptera: Lymantriinae: *Gynaephora*) endemic to the Qinghai-Tibetan Plateau"

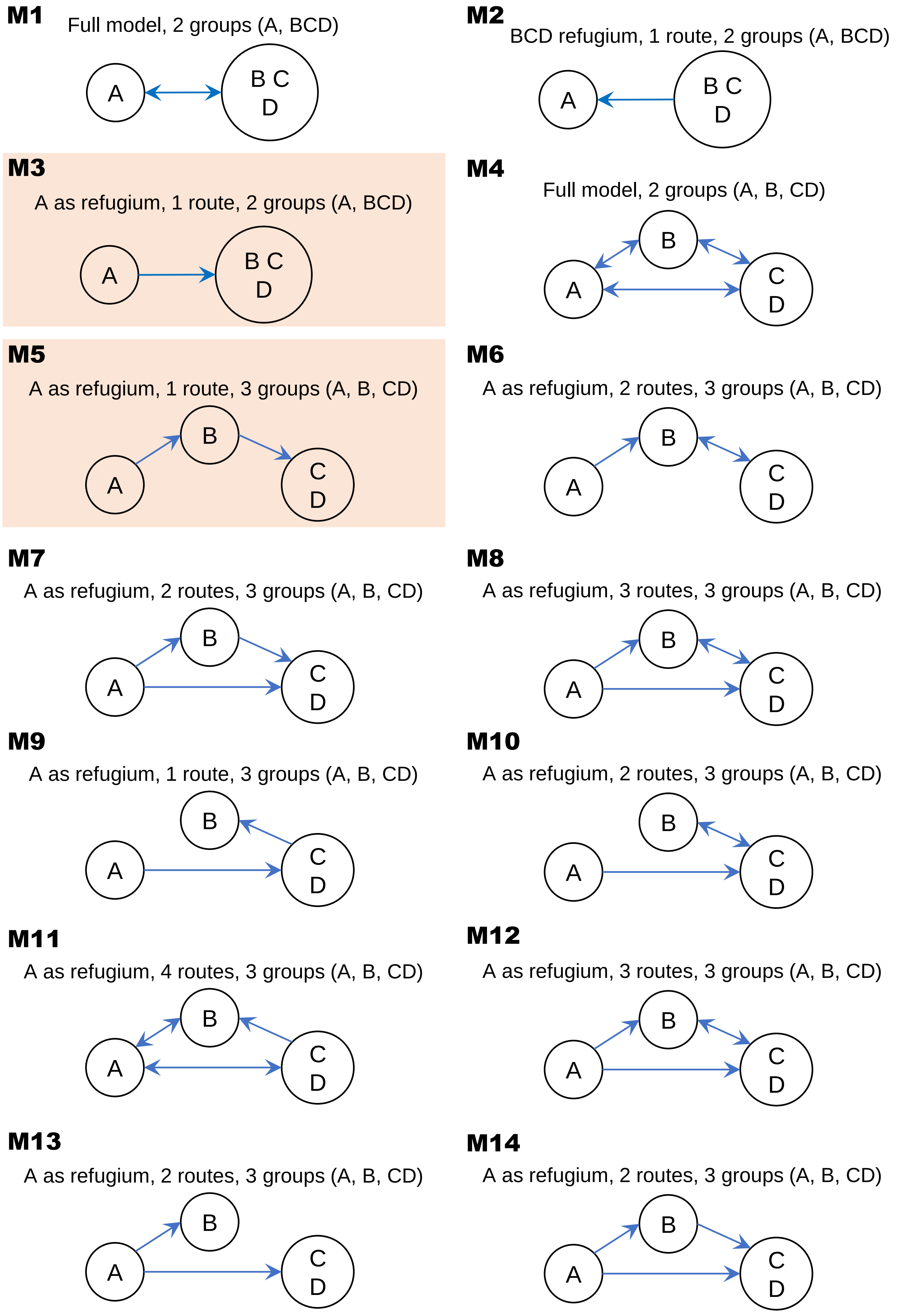
M1
Full model, 2 groups (A, BCD)
A
B C
D
M2
BCD refugium, 1 route, 2 groups (A, BCD)
A
B C
D
M3
A as refugium, 1 route, 2 groups (A, BCD)
A
B C
D
M4
Full model, 2 groups (A, B, CD)
B
A
C
D
M5
A as refugium, 1 route, 3 groups (A, B, CD)
B
A
C
D
M6
A as refugium, 2 routes, 3 groups (A, B, CD)
B
A
C
D
M7
A as refugium, 2 routes, 3 groups (A, B, CD)
B
A
C
D
M8
A as refugium, 3 routes, 3 groups (A, B, CD)
B
A
C
D
M9
A as refugium, 1 route, 3 groups (A, B, CD)
B
A
C
D
M10
A as refugium, 2 routes, 3 groups (A, B, CD)
B
A
C
D
M11
A as refugium, 4 routes, 3 groups (A, B, CD)
B
A
C
D
M12
A as refugium, 3 routes, 3 groups (A, B, CD)
B
A
C
D
M13
A as refugium, 2 routes, 3 groups (A, B, CD)
B
A
C
D
M14
A as refugium, 2 routes, 3 groups (A, B, CD)
B
A
C
D

## M15

A as refugium, 2 routes, 3 groups (A, B, CD)

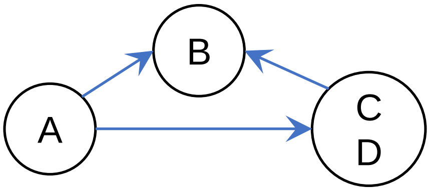


## M16

A as refugium, 3 routes, 3 groups (A, B, CD)

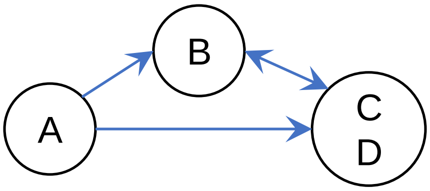


## M17

A as refugium, 2 routes, 3 groups (A, B, CD)

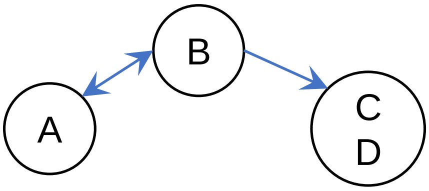


## M18

Full model, 4 groups (A, B, C, D)

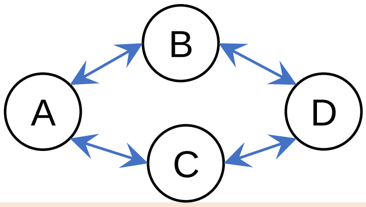


## M19

A as refugium, 1 route, 4 groups (A, B, C, D)

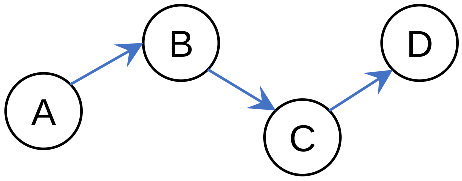


## M20

A as refugium, 2 routes, 4 groups (A, B, C, D)

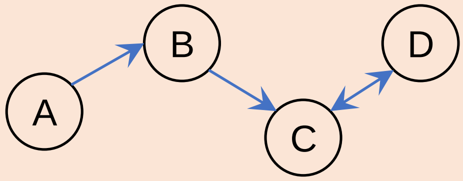


## M21

A as refugium, 1 route, 4 groups (A, B, C, D)

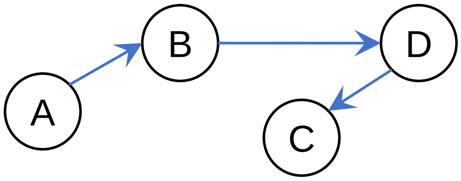


## M22

A as refugium, 2 routes, 4 groups (A, B, C, D)

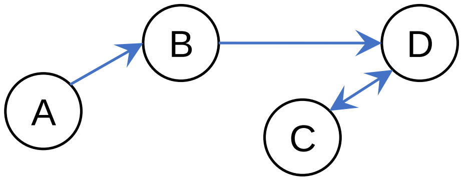


## M23

A as refugium, 2 route, 4 groups (A, B, C, D)

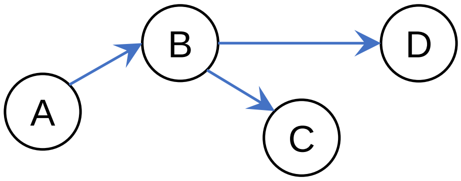


## M24

A as refugium, 3 routes, 4 groups (A, B, C, D)

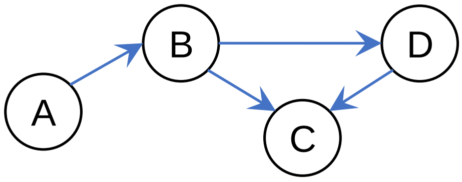


## M25

A as refugium, 2 routes, 4 groups (A, B, C, D)

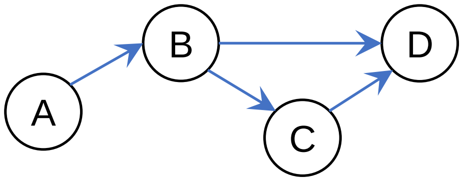


## M26

A as refugium, 3 routes, 4 groups (A, B, C, D)

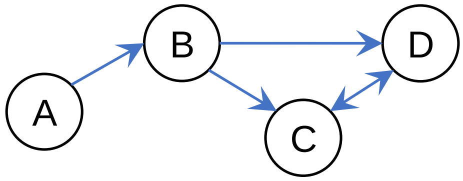
