## Supplemental Table S1 for "Mitochondrial phylogeography of grassland caterpillars (Lepidoptera: Lymantriinae: *Gynaephora*) endemic to the Qinghai-Tibetan Plateau"

**Table S1** Sampling information and haplotype distributions for 39 geographic populations of QTP *Gynaephora* species.

| Population ID | Locality | Sample size | Haplotype | Longitude | Latitude | Altitude (m) |
| --- | --- | --- | --- | --- | --- | --- |
| AB | Aba County, Sichuan Province, China | 10 | H1, H2 | 101°45'E | 32°54'N | 3350 |
| AD | Anduo County, the Tibet Autonomous Region, China | 20 | H3 | 91°68'E | 32°03'N | 4900 |
| BB | Qilian County, Qinghai Province, China | 10 | H4, H5 | 100°26'E | 38°16'N | 2800 |
| BY | Ruoergai County, Sichuan Province, China | 10 | H6, H7, H8, H9 | 103°06'E | 33°34'N | 3450 |
| CD | Chenduo County, Qinghai Province, China | 10 | H10 | 97°34'E | 33°34'N | 4000 |
| DLM | Zeku County, Qinghai Province, China | 11 | H11 | 101°55'E | 35°02'N | 3600 |
| DM | Ruoergai County, Sichuan Province, China | 10 | H6, H7, H8, H9, H12, H13 | 102°57'E | 33°18'N | 3450 |
| DR | Dari County, Qinghai Province, China | 9 | H14, H15 | 100°03'E | 33°28'N | 4150 |
| EB | Qilian County, Qinghai Province, China | 20 | H4 | 100°94'E | 37°95'N | 3400 |
| GD | Gande County, Qinghai Province, China | 9 | H16 | 100°14'E | 34°13'N | 4100 |
| GH | Luqu County, Gansu Province, China | 15 | H7, H17 | 102°36'E | 34°24'N | 3450 |
| HC | Qilian County, Qinghai Province, China | 10 | H5, H18 | 101°19'E | 37°62'N | 3200 |
| HL | Qilian County, Qinghai Province, China | 10 | H5, H18 | 100°78'E | 37°70'N | 3400 |
| HRG | Gangcha County, Qinghai Province, China | 12 | H4, H19 | 100°37'E | 37°19'N | 3400 |
| JG | Maqin County, Qinghai Province, China | 10 | H20, H21 | 100°42'E | 34°39'N | 3200 |
| KRM | Tianjun County, Qinghai Province, China | 16 | H4 | 98°84'E | 37°33'N | 3200 |
| KS | Jiuzhi County, Qinghai Province, China | 13 | H2 | 101°29'E | 33°22'N | 3650 |
| ML | Qilian County, Qinghai Province, China | 10 | H5, H30, H31, H32 | 100°44'E | 37°56'N | 3400 |
| MQ | Maqu County, Gansu Province, China | 10 | H22 | 101°52'E | 33°50'N | 3600 |
| MY | Menyuan County, Qinghai Province, China | 10 | H5 | 101°18'E | 37°38'N | 3100 |
| NG | Shiqua County, Sichuan Province, China | 20 | H23, H24, H25, H26 | 98°12'E | 32°96'N | 4400 |
| NQ | Naqu County, the Tibet Autonomous Region, China | 10 | H3 | 92°04'E | 31°48'N | 4550 |
| NR | Nierong County, the Tibet Autonomous Region, China | 16 | H3, H27, H28, H29 | 92°26'E | 31°74'N | 4700 |
| QML | Qumalai County, Qinghai Province, China | 9 | H30, H31, H32 | 95°79'E | 34°07'N | 4550 |
| RW | Suoxian County, the Tibet Autonomous Region, China | 20 | H3 | 93°82'E | 31°72'N | 3600 |
| SD | Seda County, Sichuan Province, China | 20 | H33 | 100°33'E | 32°27'N | 4000 |
| SHRM | Jiuzhi County, Qinghai Province, China | 10 | H33 | 101°19'E | 33°22'N | 4050 |
| SJC | Gangcha County, Qinghai Province, China | 16 | H4, H5, H34 | 100°16'E | 37°35'N | 3400 |
| SS | Anduo County, the Tibet Autonomous Region, China | 18 | H3 | 92°05'E | 31°93'N | 4800 |
| TD | Tongde County, Qinghai Province, China | 10 | H35 | 100°48'E | 34°42'N | 3400 |
| TR | Tongren County, Qinghai Province, China | 10 | H7, H11 | 101°47'E | 35°12'N | 3500 |
| XH | Haiyan County, Qinghai Province, China | 12 | H4 | 100°91'E | 37°03'N | 3050 |
| XQ | Biru County, the Tibet Autonomous Region, China | 20 | H3 | 93°83'E | 31°79'N | 4400 |
| YGN | Henan County, Qinghai Province, China | 13 | H11, H36, H37 | 101°36'E | 34°46'N | 3700 |
| YS | Yushu County, Qinghai Province, China | 10 | H38 | 97°13'E | 32°83'N | 4000 |
| ZD | Zhiduo County, Qinghai Province, China | 10 | H30, H39 | 95°46'E | 33°79'N | 4500 |
| ZK | Zeku County, Qinghai Province, China | 10 | H11, H40 | 101°31'E | 34°54'N | 3650 |
| ZQ | Zaduo County, Qinghai Province, China | 8 | H7, H11, H30 | 95°21'E | 33°22'N | 4250 |
| ZQZ | Zeku County, Qinghai Province, China | 11 | H7, H11 | 101°45'E | 35°00'N | 3650 |
