## Supplemental Table S3 for "Mitochondrial phylogeography of grassland caterpillars (Lepidoptera: Lymantriinae: *Gynaephora*) endemic to the Qinghai-Tibetan Plateau"

**Table S3** Mantel tests for relationships between genetic and geographic distances of the QTP *Gynaephora* species. See Figure 1 for Clade A, Clade B, Clade C, and Clade D.

|  | Genetic distance | Geographic distance  (GD) | Correlation  coefficient (*r*) | *P* value |
| --- | --- | --- | --- | --- |
| All 39 populations | *F*_ST_ | GD | 0.373 | <0.01 |
|  | Log (*F*_ST_) | GD | 0.169 | <0.01 |
|  | *F*_ST_ | Ln GD | 0.517 | <0.01 |
|  | Log (*F*_ST_) | Ln GD | 0.222 | <0.01 |
| Clade A | *F*_ST_ | GD | 0.457 | <0.01 |
|  | Log (*F*_ST_) | GD | 0.191 | <0.05 |
|  | *F*_ST_ | Ln GD | 0.191 | <0.05 |
|  | Log (*F*_ST_) | Ln GD | 0.356 | <0.05 |
| Clade B | *F*_ST_ | GD | -0.247 | 0.837 |
|  | Log (*F*_ST_) | GD | -0.284 | 0.847 |
|  | *F*_ST_ | LnGD | -0.096 | 0.583 |
|  | Log (*F*_ST_) | LnGD | -0.132 | 0.551 |
| Clade C | *F*_ST_ | GD | -0.094 | 0.713 |
|  | Log (*F*_ST_) | GD | 0.020 | 0.516 |
|  | *F*_ST_ | LnGD | 0.017 | 0.418 |
|  | Log (*F*_ST_) | LnGD | 0.004 | 0.461 |
| Clade D | *F*_ST_ | GD | 0.149 | 0.168 |
|  | Log (*F*_ST_) | GD | 0.201 | 0.086 |
|  | *F*_ST_ | LnGD | 0.175 | 0.138 |
|  | Log (*F*_ST_) | LnGD | 0.206 | 0.085 |
