## Supplemental Table S4 for "Mitochondrial phylogeography of grassland caterpillars (Lepidoptera: Lymantriinae: *Gynaephora*) endemic to the Qinghai-Tibetan Plateau"

**Table S4** Parameter estimates of Theta (θ) and immigration rate (M) for migration model comparisons based on two mitochondrial genes (*cox1* and *nad5*). Mean values and 2.5–97.5% intervals in brackets are provided for each parameter. Detailed information for the three best models (M3, M5, and M20) is provided in Table 3 and Figure S2.

| Model | Group | θ1 | θ2 | θ3 | θ4 | M_1->2_ | M_2->3_ | M_3->4_ | M_4->3_ |
| --- | --- | --- | --- | --- | --- | --- | --- | --- | --- |
| M20 | A group (1)  B group (2)  C group (3)  D group (4) | 0.00562  (0.00153~0.00973) | 0.00110  (0~0.00293) | 0.00089  (0~0.00267) | 0.00213  (0~0.00413) | 91.2  (2.0~214.7) | 142.9  (0~380.0) | 104.4  (0~268.7) | 177.6  (0~503.3) |
| M3 | A group (1)  BCD group (2) | 0.00804  (0~0.00467) | 0.00488  (0.00193~0.00773) | – | – | 31.2  (0~94.7) | – | – | – |
| M5 | A group (1)  B group (2)  CD group (3) | 0.00858  (0.00487~0.01320) | 0.00110  (0~0.00293) | 0.00292  (0.00053~0.00520) | – | 118.3  (8.0~268.0) | 76.5  (0~171.3) | – | – |
